# Genomics of sex allocation in the parasitoid wasp *Nasonia vitripennis*

**DOI:** 10.1101/2020.03.18.997619

**Authors:** Bart A. Pannebakker, Nicola Cook, Joost van den Heuvel, Louis van de Zande, David M. Shuker

## Abstract

**Background:** Whilst adaptive facultative sex allocation has been widely studied at the phenotypic level across a broad range of organisms, we still know remarkably little about its genetic architecture. Here, we explore the genome-wide basis of sex ratio variation in the parasitoid wasp *Nasonia vitripennis*, perhaps the best studied organism in terms of sex allocation, and well known for its response to local mate competition (LMC).

**Results:** We performed a genome-wide association study (GWAS) for single foundress sex ratios using iso-female lines derived from the recently developed outbred *N. vitripennis* laboratory strain HVRx. The iso-female lines capture a sample of the genetic variation in HVRx and we present them as the first iteration of the *Nasonia vitripennis* Genome Reference Panel (NVGRP 1.0). This panel provides an assessment of the standing genetic variation for sex ratio in the study population. Using the NVGRP, we discovered a cluster of 18 linked SNPs, encompassing 9 annotated loci associated with sex ratio variation. Furthermore, we found evidence that sex ratio has a shared genetic basis with clutch size on three different chromosomes.

**Conclusions:** Our approach provides a thorough description of the quantitative genetic basis of sex ratio variation in *Nasonia* at the genome level and reveals a number of inter-related candidate loci underlying sex allocation regulation.

## Background

The study of sex allocation is one of the most successful areas in evolutionary biology [1–3]. Theoretical predictions of optimal resource allocation to male and female offspring in response to environmental conditions are now supported by a wealth of empirical data [1, 4–9]. This is particularly true for Local Mate Competition (LMC) theory [10, 11]. Briefly, LMC theory describes sex allocation dynamics when related males (including full siblings) compete for mates in locally structured populations. When kin competition for mates is high, termed local mate competition by Hamilton (1967), female-biased sex ratios, that limit kin competition for mates and maximise the number of available mates for those competing males, are favoured [12]. LMC is maximal when the male offspring from a single female compete for mates (their sisters) on a discrete mating patch. When LMC is reduced, with increasing numbers of unrelated males competing for mates, less female-biased sex ratios are favoured. The logic of Hamilton’s LMC theory extends to situations where females experience environments that vary in the expected level of LMC amongst sons, for instance when different numbers of females oviposit together on a patch. Facultative sex allocation is then predicted, and has been shown in a diverse range of organisms, from malarial parasites to fig wasps [3, 13]. Perhaps the most extensive exploration of LMC theory has occurred in parasitoid wasps, however, especially *Nasonia vitripennis* [14, 15].

Despite the success of sex allocation theory at the phenotypic level, we still know rather little about the underlying mechanisms of sex allocation at the molecular level [16–19]. This is important for at least three reasons. First, deviations in sex ratio from theoretical predictions still remain [20] and require explanation. In *Nasonia vitripennis*, extending and testing basic LMC theory has revealed that understanding how female parasitoids obtain and use information relating to the expected level of LMC their sons will experience can refine theoretical models and explain much of this variation [15, 21–25]. However, there is also LMC-independent genetic variation in sex allocation in *N*. *vitripennis* [19, 26–30]. Second, adaptive sex allocation provides an opportunity to explore the mechanisms underlying adaptation, including the genetic architecture and molecular evolution of a well-characterised trait (for recent discussions see for example; [31–36]). For instance, to what extent does genetic architecture constrain sex allocation? Third, phenotypic plasticity is at the heart of facultative sex allocation, and there has long been interest in how such plasticity is encoded in the genome [37]. Altogether, sex allocation is a well-characterised plastic trait, offering the opportunity to dissect the genetic basis of plasticity.

To date, studies on the genetic basis of sex ratio have focussed on quantitative genetic analysis across a range of organisms, including *Drosophila* [38, 39], ants [40], snails [41], fish[42], turtles [43], birds [44], pigs [45] and humans [46]. Several species of parasitoid wasp also show genetic variation for sex ratio [47–51], but most work has focused on *Nasonia vitripennis* [26–28, 30, 52]. More recently, mutation accumulation studies have explored how mutation augments additive genetic variation in sex ratio [16], suggesting that genes governing sex ratio are pleiotropic, also influencing other fitness-related traits. A linkage mapping study of sex ratio variation within a natural population of *N. vitripennis* identified a major quantitative trait locus (QTL) on chromosome 2 and three weaker QTL, one on chromosome 3 and two on chromosome 5. In addition, the QTL data revealed scope for pleiotropy between offspring sex ratio and clutch size [19]. Finally, recent work has explored patterns of gene expression associated with sex allocation in *Nasonia*. Thus far, patterns of differential gene expression associated with oviposition behaviour have been revealed in females, but there is no suggestion that the facultative response to LMC cues made during oviposition is associated with any change in gene expression [17, 18, 53]. This suggests that, whilst oviposition is associated with changes in gene expression, phenotypically plastic sex allocation is not.

Here, in order to better understand the genomics of facultative sex allocation and oviposition, we extend our characterisation of the genetic basis of sex ratio variation in *N*. *vitripennis* with a genome-wide association study (GWAS) for single foundress sex ratios using iso-female lines derived from the recently developed outbred laboratory strain HVRx [54]. The iso-female lines capture a sample of the genetic variation in HVRx and we present them as the first iteration of the *Nasonia vitripennis* Genome Reference Panel (NVGRP 1.0). This panel provides an assessment of the standing genetic variation for sex ratio in the study population. Our approach provides a thorough description of the quantitative genetic basis of sex ratio variation in *Nasonia* and reveals a number of candidate molecular processes underlying sex allocation.

## Results

### Molecular variation in NVGRP

The NVGRP was constructed by isolating mated females from the HVRx outbred laboratory population, followed by 9 generations of inbreeding. 34 NVGRP lines were sequenced using Illumina HiSeq 2000, and the reads were mapped to the *N. vitripennis* reference genome. The mean sequence depth was 5.4X per line, with on average 230.2 Mb (96.47%) of the effective reference sequence covered per line (Supplementary Table 1). Using JGIL [55], which takes into account the expected allele frequencies after 9 generations of inbreeding from the outbred HVRx laboratory population, we called 205,691 SNPs. The residual heterozygosity in the NVGRP lines was low, on average the proportion of segregating SNPs across lines was 2.68% (s.e.=0.35%), which is less than is theoretically expected after 9 generations of inbreeding (13% given an inbreeding coefficient of *F*=0.87). The majority of the SNPs in the NVGRP (96,920) are found in intergenic regions (Figure 1). 6,250 SNPs occur in the 5’ UTR and 7086 in the 3’ UTR sequences. In the coding regions, there are 12,344 synonymous SNPs and 6,723 nonsynonymous SNPs (6,634 missense and 89 nonsense SNPs), resulting in a *d*_*N*_*/d*_*S*_ ratio of 0.545.

**Figure 1.**
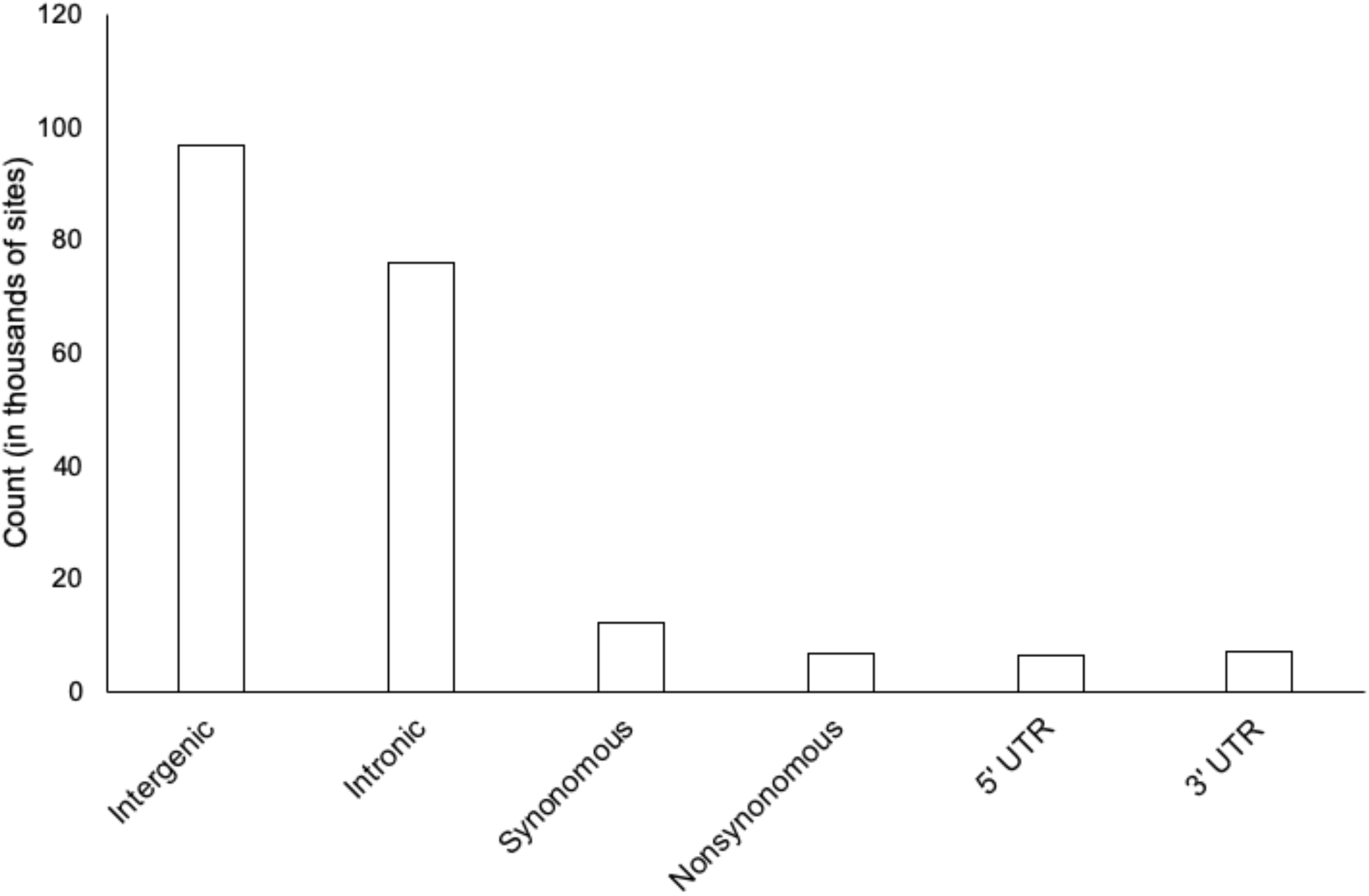
SNP counts per site class. SNPs were attributed to only one class, according to the SnpEff reporting order (most severe effects first; see [114]).

### NVGRP population genetics

We calculated the genome-wide polymorphism (Watterson’s theta (*θ*) and nucleotide diversity (*π*)), both per chromosome, and in 400 kb non-overlapping windows (Figure 2, Supplementary Table 2). Averaged over the entire genome, *π* = 1.204e-03 (s.e.= 2.841e-05), and *θ* = 1.208e-03 (s.e.= 2.823e-05). The average polymorphism differs per chromosome for both *π* (*F*_*4, 577*_=3.047, *P*=0.017) and *θ* (*F*_*4, 577*_=2.903, *P*=0.021). Nucleotide diversity is higher towards the tips of the chromosomes (Figure 2). Comparison of the NVGRP to the HVRx source population shows similar levels of the nucleotide diversity (*π*), indicating the 34 NVGRP to be a good representation of the variation present in the source population. However, Watterson’s theta (*θ*) is lower over all the NVGRP lines compared to HVRx, suggesting a loss of low frequency alleles, which is further reflected in the lower genome-wide Tajima’s D values (Figure 2). Linkage disequilibrium (LD) decays to half the of the initial value in 17.8 kb (half-decay distance, Figure 3), with an average pairwise *r*^*2*^ value of 0.08 (s.e.=0.0005). The extend of LD varies across and within the chromosomes. Some regions on chromosomes 1 and 3 exhibit half-decay distances over 8.9Mb and 36Mb respectively (Figure 4). Pairwise *F*_*ST*_ values between the NVGRP lines were on average 0.309 (s.d. = 0.046) with a minimum of 0.145 and a maximum of 0.465. This indicates that inbreeding on the one hand has resulted in highly divergent lines, but, on the other hand, ample overlap in genetic variation still exists between all pairs of lines (i.e., we did not find lines completely divergent for all SNPs). Furthermore, the pairwise *F*_*ST*_ values did not correlate with the absolute differences in sex ratio between lines selected for the GWAS (Spearman’s *r*_s_= −0.034, *P*=0.542) indicating that no correction for population structure in the GWAS is needed.

**Figure 2.**
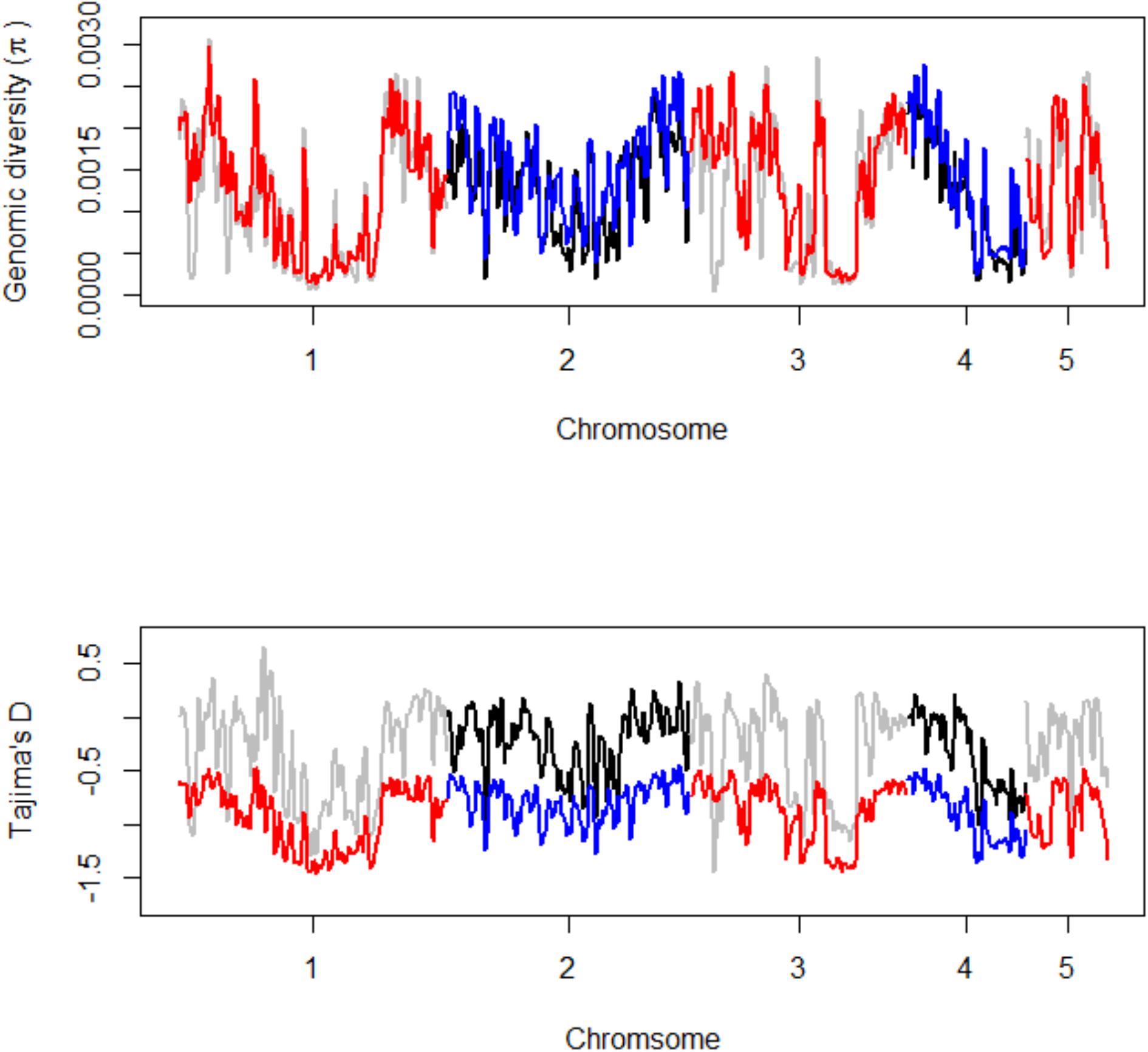
Mean nucleotide diversity *π* (upper panel) and Tajima’s D (lower panel) over 400 kb windows across the chromosomes in NVGRP (red and blue lines) and the HVRx laboratory outbred population (grey and black lines).

**Figure 3.**
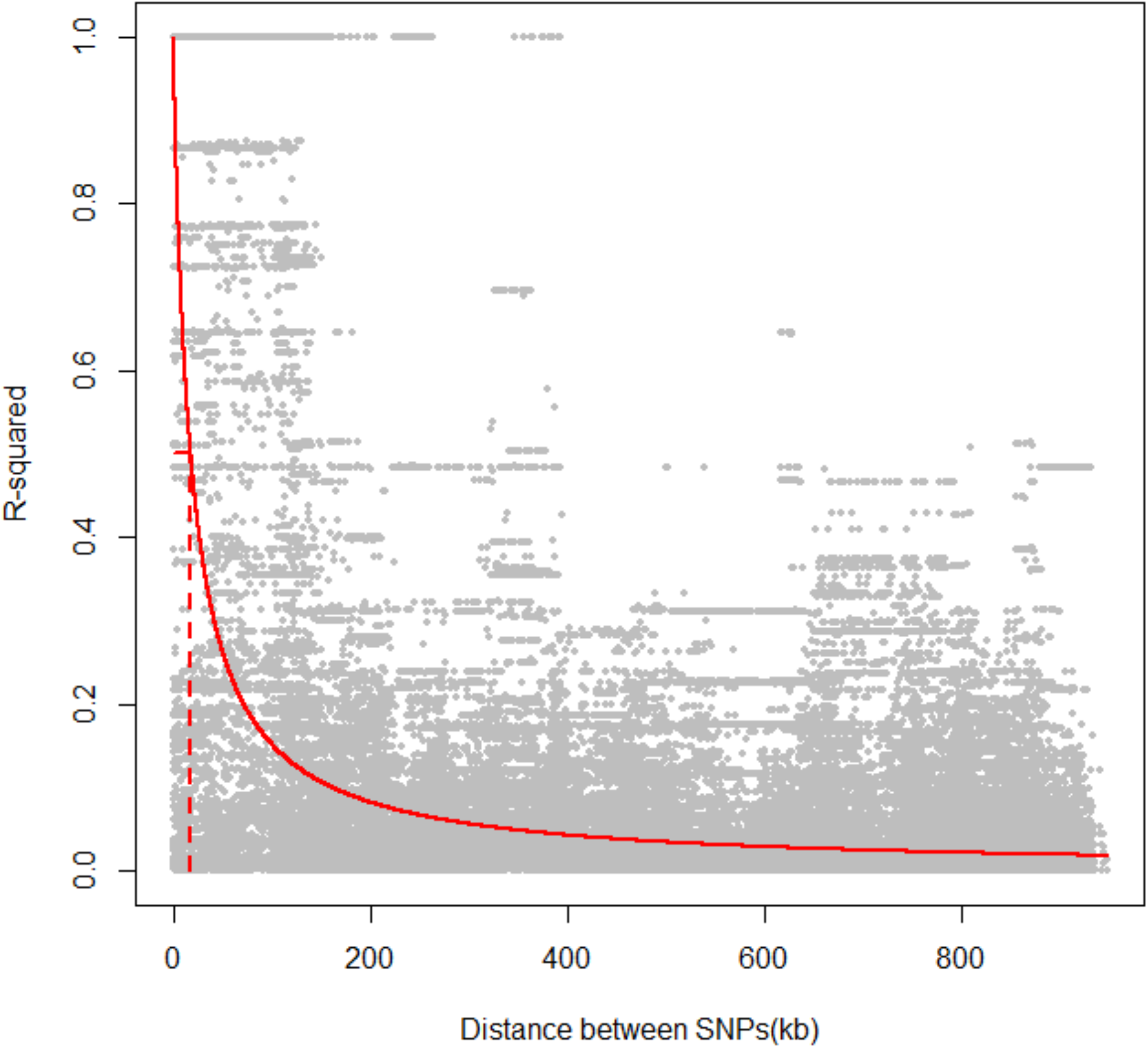
Decay of linkage disequilibrium with physical distance. Dots shows the *r*^*2*^ among pairs of SNPs, red solid line gives the non-linear least squares fit of *r*^*2*^ on the distance between pairs of SNP. Dashed line indicates the half-decay LD distance at 17.8 kb.

**Figure 4.**
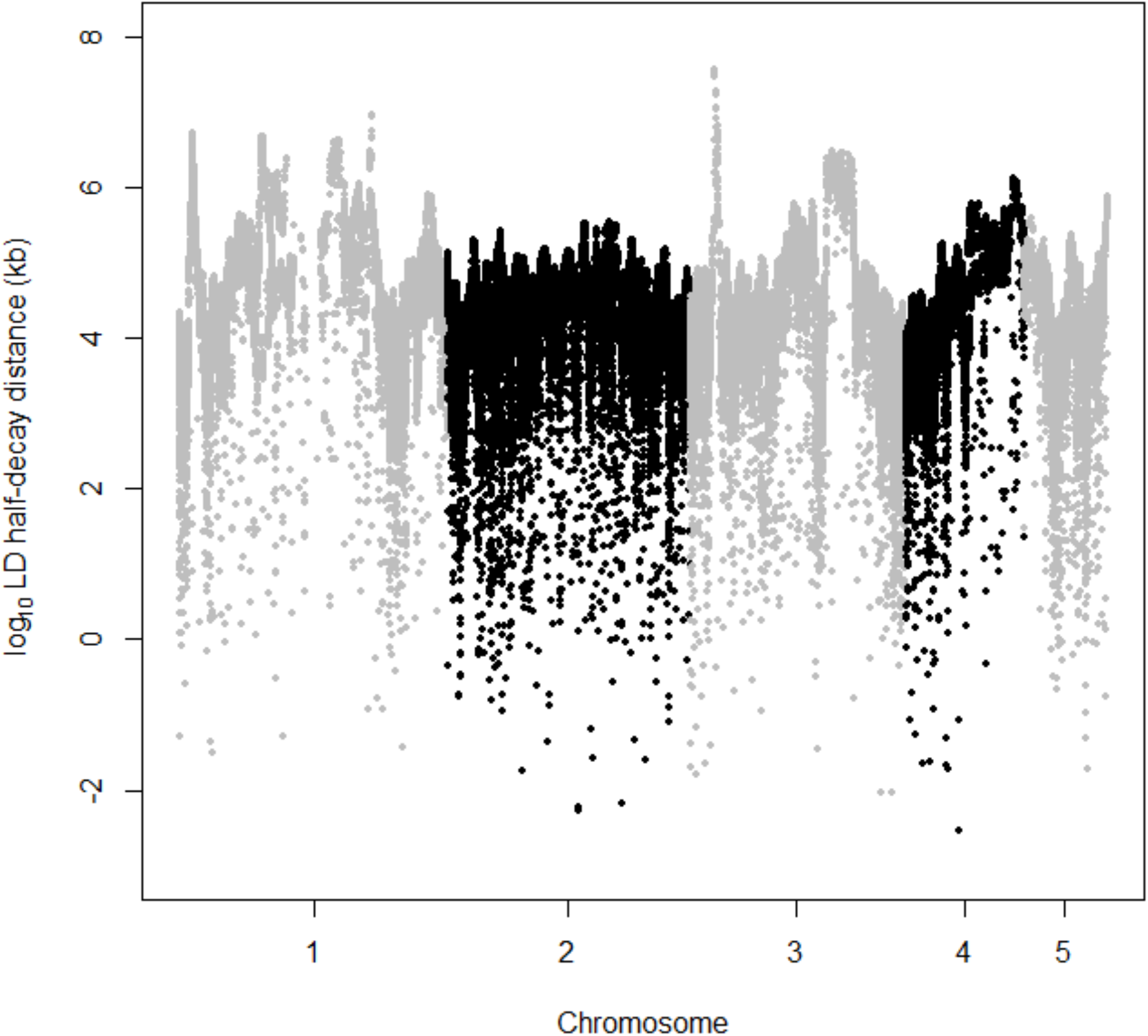
Linkage disequilibrium half-decay distance across the genome. For each SNP on each of the five chromosomes, the log_10_(LD half-decay distance) is plotted.

### Sex ratio GWAS

To exclude possible artefacts owing to differences in *Wolbachia* infection status, we only collected sex ratio data from NVGRP lines testing positive for *Wolbachia*-infection (Supplementary Figure 1). Among these, line 62 was a clear outlier, showing very male-biased sex ratios and low clutch sizes, indicative of a transient *Wolbachia* infection status (Supplementary Figure 2) resulting in within-line cytoplasmic incompatibility. We therefore excluded line 62 from further analysis. Across these 25 NVGRP lines, there was highly significant among-line variation in sex ratio (Binomial GLM: *F*_*24,918*_ = 15.244, *P*<0.0001). This among-line variation represents a broad-sense heritability of *H*^*2*^ = 0.106 for sex ratio (Likelihood Ratio=56.176, *P*<0.0001). For clutch size, the 25 NVGRP lines showed a broad-sense heritability of *H*^*2*^ = 0.078 (Likelihood Ratio=36.61, *P*<0.0001). Among the NVGRP lines, our data showed a weak, non-significant negative correlation between sex ratio and clutch size (Supplementary Figure 3, linear regression: *b* = −0.0015 (s.e.= 0.0008), *t*=−1.823, *P*=0.08).

In total, 18 SNPs were significantly associated with sex ratio, according to our empirical FDR threshold of 0.1. (Figure 5, Table 1, Supplementary Table 3). These SNPs represent a single peak on chromosome 1 and show a high degree of linkage (mean *r*^*2*^=0.97, Supplementary Table 4). All SNPs represent common variants, with an average minimal allele frequency (MAF) of 0.416. These 18 SNPs were associated with 9 annotated loci (Table 1). While the majority of the significant SNPs are predicted to either be intron variants (8) or upstream variants (7) of genes, we found two significant SNPs with a potential effect on the expression of the associated loci. NC_015867.2_30570243 is causing a synonymous change (p.Ala224Ala) in the splice region of exon 4 of *cilia- and flagella-associated protein 52* (CFAP52, LOC100124028), while NC_015867.2_30604770 (*P*= 5.00^*^10^−7^) is a variant in the 5’ untranslated region (UTR) of *ADP-ribosylation factor-like protein 4C* (Arl4C, LOC100124030). We will discuss both these SNPs in more detail below.

**Figure 5.**
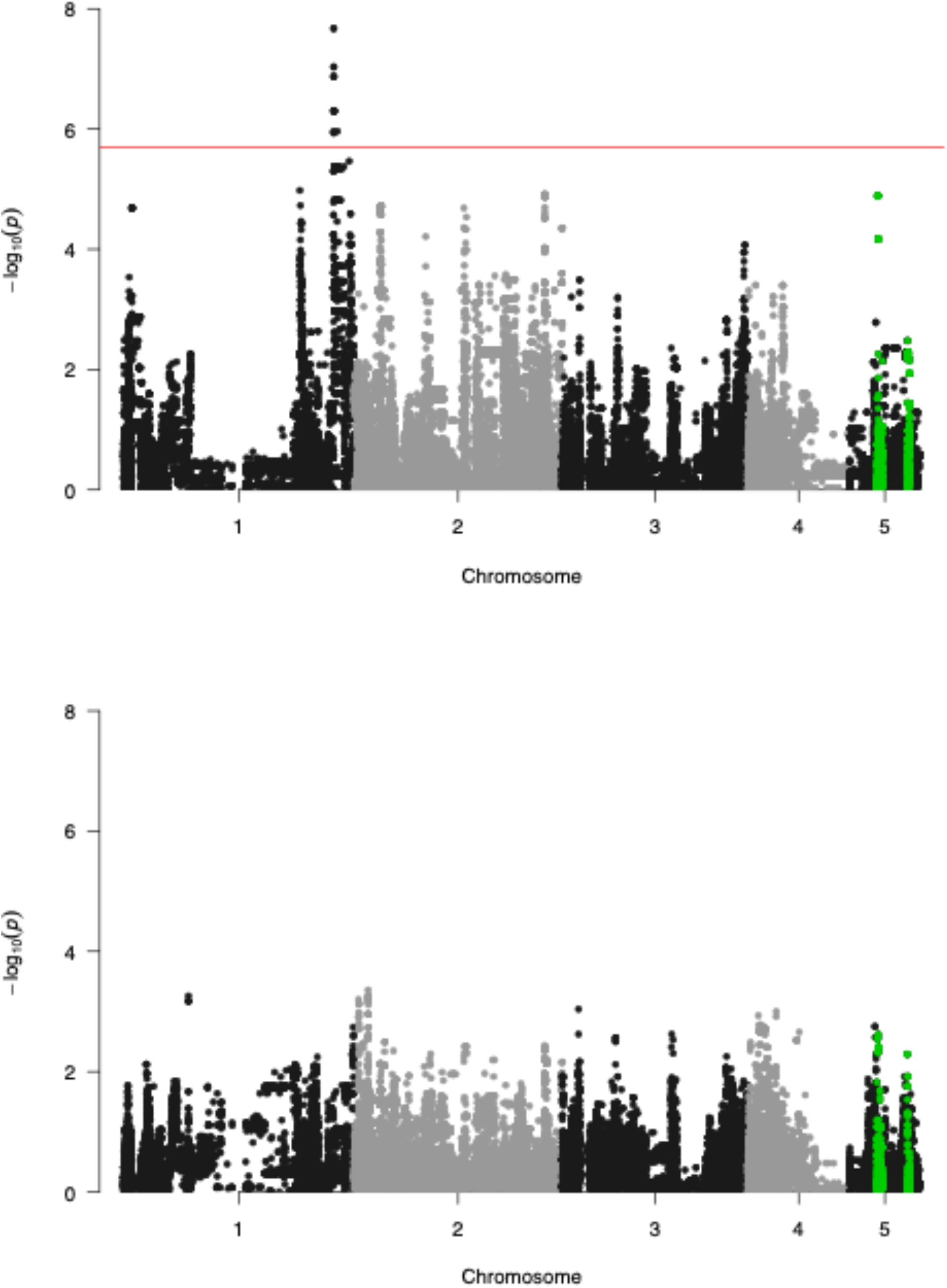
Manhattan plot for offspring sex ratio (top panel) and brood size (bottom panel) in *N. vitripennis* in the GWAS experiment, showing −log10(P)-values of the single marker regressions for every polymorphic SNP across the chromosomes of the *N. vitripennis* Genetic Reference Panel (NVGRP). Red line indicates the empirical threshold at a q-value of 0.1, corresponding to *P=2.00*^*\**^*10*^*−6*^, *or −log*_*10*_*(P)*= 5.70 for sex ratio. Green highlighted SNPs show the 400kb windows in which the P-values for sex ratio and brood size overlap more than expected by chance.

We found no SNPs significantly associated with clutch size variation across the lines. However, despite this absence, we did detect significant enrichment of overlap between the *P*-values of the sex ratio and clutch size GWAS in 400kb windows on chromosomes 2, 3 and 5 (Supplementary Figure 4). Most notably, two regions on chromosome 5, between 4,000kb – 4,800kb and between 8,400kb – 8,800kb show enriched overlap in test statistics, indicating a shared genetic background for these two traits (Figure 5). Although our finding of 18 strongly linked variants in the same genomic region suggests that the number of variants in the NVGRP segregating for sex ratio is limited, this overlapping background of sex ratio with clutch size on three different chromosomes, in addition to the a large number of sub-significant peaks for sex ratio throughout the genome, suggests that sex allocation is a complex trait with polygenic regulation and epistatic interactions.

## Discussion

Sex allocation has been extensively studied at the phenotypic level, especially in species such as parasitoid wasps. However, our understanding of the genetic architecture of the mechanisms associated with sex allocation is much more rudimentary. Here we have shown that at least 18 single nucleotide polymorphisms are associated with variation in sex allocation across experimental lines drawn from a single population. While only a single peak significantly passed the genome-wide threshold, both the presence of a large number of subsignificant peaks (Figure 5), and evidence of an overlapping genetic background on chromosome 5, confirm the polygenic nature of the genetic variation in sex allocation in *Nasonia*. While the emerging picture is that variation in most complex traits is typically influenced by a large number of loci with pleiotropic alleles, polygenic traits represent a major challenge in terms of understanding the molecular genetic basis of phenotypes and their adaptive evolution [36]. This is the case in terms of both understanding how interactions among DNA sequences constrain or promote the production of adaptive phenotypes, and also in terms of reconstructing the path evolution has taken at the molecular level, including identifying key substitutions. Put simply, we are likely to have an embarrassment of riches, although even then, we may well only be able to discover a fraction of evolutionarily relevant SNPs [32]. Moreover, unlike candidate gene approaches, we are often faced with genes or putative regulatory sequences about which we know little or nothing, either through a lack of annotation, or through the lack of a clear link between proposed gene ontology and the phenotype of interest. In our case, the 18 SNPs on chromosome 1 that are associated with sex ratio variation among our lines, have predicted effects on 9 annotated loci.

Of these, two significant SNPs are predicted to have an effect on the expression of the loci they are associated with. NC_015867.2_30570243 (*P*=1.12^*^10^−6^) is a variant causing a synonymous change (p.Ala224Ala) in the splice region of exon 4 of *cilia- and flagella-associated protein 52* (CFAP52, LOC100124028). CFAP52 is highly conserved and its human ortholog, WDR16, occurs in sperm tails [56]. Furthermore, human low-motility sperm shows low expression of this testis specific transcript WDR16 [57]. While no direct studies have been done into the role of CFAP52 in *Nasonia* sperm, CFAP52 shows high expression levels in *N. vitripennis* testis [58, 59]. Because of its role in sperm motility, splice region changes could affect the functionality of CFAP52 and therefore directly impact sex allocation in the haplodiploid *N. vitripennis,* in which fertilization is the key trigger to female development [60]. Results such as these would help explain the small influence of males in sex allocation observed in *N. vitripennis* [61]. The observed variation among strains in sex ratio could well be due to differences in male ejaculate quality and fertilization success. Such variation in male fertility might also explain why the observed single foundress sex ratios are slightly higher (i.e. more males) than we might expect from the number of females a single male can inseminate, and would act as a form of ‘fertility insurance’ [3, 62]. Whether the splice region variant CFAP52 observed in the NVGRP can account for such variation requires a more thorough characterization of the potential mis-splicing and is outwith the scope of the present study [63].

The other SNP that may well influence gene expression is NC_015867.2_30604770 (*P*= 5.00^*^10^−7^). This is a variant in the 5’ untranslated region (UTR) of *ADP-ribosylation factor-like protein 4C* (Arl4C, LOC100124030), but does not generate a premature start codon. The 5’ UTR is the mRNA region directly upstream from the start codon and plays a role in the regulation of transcript translation. While it is hard to predict the exact effect of a 5’ UTR variant in a non-model system, 5’ UTR variants can affect the translation efficiency and hence the expression level of proteins [64]. Arl4 are group of proteins that belongs to the family of ADP-ribosylation factors, which are involved in the regulation of vesicular trafficking processes [65]. Arl4C has been implicated in the regulation of vesicular traffic of cholesterol and of transferrin in endosomes [66] and likely plays a similar role as the related Arl4A [67]. Interestingly, Arl4A is strongly expressed in adult testis in mice [68] and targeted disruption of Arl4A resulted in a reduced sperm count [69]. More functional studies are needed to determine if Arl4C also plays a role in *Nasonia* spermatogenesis, and whether the identified 5’ UTR variant affects Arl4C expression and alters the *Nasonia* sex ratio phenotype, again via effects on sperm quality or quantity.

While no significant SNPs were found for clutch size, we did detect significant enrichment of overlap between the *P*-values of the sex ratio and clutch size GWAS in 400kb windows on chromosomes 2, 3 and 5, with those on chromosome 5 providing the strongest evidence (i.e. *P*-values were non-randomly associated with each other for the two traits across these regions, whereas there should be no association if there is no shared genetic basis: Figure 5, Supplementary Figure 4). This overlap indicates a shared genetic basis for the two traits, compatible with theoretical predictions on the genetic basis of sex ratio. Based on the observed natural variation and estimates of mutational parameters for sex ratio, the genetic variation for sex ratio is predicted to be maintained by selection on pleiotropic loci with effects on other fitness related traits [16]. While clutch size is only one of many fitness-related traits that can show pleiotropy with sex ratio, a correlation between these two traits is also expected from theoretical work. In the case of a single foundress per host, LMC theory predicts that a mother should only produce enough sons to mate with all of her daughters [10]. For small clutch sizes, this can be a single male, and when clutch size increases, an increasingly male-biased sex ratio is predicted [20, 70, 71]. Interestingly, our previous analysis of the quantitative genetic basis of sex ratio and clutch size in different Dutch *N. vitripennis* strains, also identified overlapping QTLs for both traits on chromosomes 2 and 5 on a recombination linkage map [72]. Unfortunately, of the eight microsatellite markers associated to the sex ratio in the QTL study, only two could be found on the same chromosomes in the current assembly (Nvit_2.1). The other six are located on unlocalized scaffolds (data not shown), making a direct comparison between these studies difficult and indicating the need for a further improved genome assembly for this species [73].

In addition to revealing some of the molecular genetic variants associated with sex allocation in *Nasonia*, we have also presented a new genomic resource for the community. Since the publication of the *Nasonia* genome in 2010 [74], there has been a steadily growing number of studies of the molecular genetics of a range of phenotypes, deploying a number of techniques [73, 75–81]. In terms of our own work on sex allocation, we have shown for example that facultative sex allocation under LMC is not associated with changes in gene expression [17, 18], even though female oviposition of eggs is associated with major changes to the transcriptome (in particular, a down-regulation of metabolic processes: [17, 53]). We have also shown though that disrupting patterns of DNA methylation changes the pattern of facultative sex allocation [82, 83], suggesting that the regulation of gene expression via DNA methylation is important for facultative sex allocation.

Our first iteration of a proposed *Nasonia vitripennis* Genome Reference Panel (NVGRP 1.0) currently consists of 34 iso-female lines generated from wasps collected from one population, of which 25 lines comprised the GWAS for sex ratio presented here. Whilst the number of lines is currently modest, we have nonetheless captured SNPs associated with sex ratio; moreover, the NVGRP lines exhibit significant broad-sense heritabilities for a range of traits, not just sex ratio, including total lifetime fecundity, longevity, head width, wing length, and starvation resistance (*H*^2^ = 0.13−0.58; B.A. Pannebakker, unpublished observations). Importantly, as a species characterised by sib-mating, and thus inbreeding, the creation of iso-female lines in *Nasonia* does not suffer the problems associated with the inbreeding of natural out-crossers [84], and so we do not see great reductions in fitness in our inbred lines. The effect of sib-mating is reflected in the relatively slow decay of linkage disequilibrium (*r*^*2*^<0.1 at 160.2kb, and *r*^*2*^<0.2 at 71.2kb) as is also observed in selfing plants (e.g. rice *Oriza sativa*, *r*^*2*^<0.1 at 75-150kb [85] or soy *Glycine max r*^*2*^<0.1 at 90-500kb [86]). Other insects, such as *Drosophila melanogaster* and *Apis mellifera*, show much shorter distances at which LD decays (*r*^*2*^<0.2 at 10 bp and *r*^*2*^<0.2 at 1kb respectively, [87, 88]).

The extent of LD in the NVGRP lines does show large variation within and across chromosomes, which likely reflects the observed variation in recombination rates in *Nasonia* [89–91]. In direct observations of recombination events from markers segregating in a cross, areas of low recombination were observed near the centre of the chromosomes [89, 90]. The NVGRP shows areas of high LD, but not just at chromosome centres. The LD estimates in the NVGRP, however, are population estimates, integrating historical recombination events that could result in a different pattern than from the present-day recombination estimates observed in the progeny from a cross [92]. To what extent natural inbreeding, or other demographic factors such as the limited population size of NVGRP, is driving the slow decay of linkage disequilibrium and the observed variation in LD in *Nasonia*, requires further genotyping of additional samples from wild populations.

## Conclusions

We present the first iteration of the *Nasonia vitripennis* Genetic Reference Panel as a community resource for the analysis of complex traits. We found substantial variation for sex allocation in the NVGRP and identified 18 SNPs associated with variation in sex allocation and found evidence for overlapping genetic background of sex ratio with clutch size on different chromosomes. As our data represent only a sample of the standing genetic variation in our study population, it is likely that we have missed other variants that influence sex ratio variation segregating in that population, but we have nonetheless provided the first genomic visualization of the heritability of sex ratio observed in this and other studies on *Nasonia* [16, 26, 28].

Wild *N. vitripennis* populations show rather limited population genetic structure across Europe, which perhaps explains our ability to capture significant genetic variation with only a comparatively small sample of lines from one population [93, 94]. Nevertheless, we recognise that the study presented here is as much a proof-of-principle for exploring the molecular genetic basis of sex allocation in *Nasonia* as anything else, and an expansion of the NVGRP is certainly required. However, we note that as whole-genome sequencing becomes ever cheaper and alternative genotyping methods are developed (such as RAD sequencing and other genotype-by-sequencing techniques [95, 96]), the role of reference panels may change. Shifting from the main focus of a study, they can provide supporting genomic resources, or genetic hypotheses to test, for studies that interrogate genetic variants in the wild more directly [97–100].

## Methods

### Study Organism

*Nasonia vitripennis* (Hymenoptera, Chalcidoidea) is a generalist parasitoid of large dipteran pupae, including species of Calliphoridae. Depending on host species, females oviposit between 20-50 eggs in an individual host, with male offspring emerging just before females (after approximately 14 days at 25°C; [101]). Male individuals are brachypterous and are unable to fly, remaining close to the emergence site where they compete with each other for emerging females, including their sisters. Females disperse after mating to locate new hosts. As with all Hymenoptera, *N. vitripennis* is haplodiploid, with diploid females developing from fertilised eggs, and haploid males developing from unfertilised eggs. Sex allocation is therefore associated with females controlling the fertilisation of eggs during oviposition, releasing stored sperm from the spermatheca to fertilise eggs as they pass down the oviduct. Unless otherwise specified, wasps were reared on *Calliphora* spp. hosts at 25°C, 16L:8D light conditions.

### Base population

We used the HVRx outbred population of *N. vitripennis* [54] as the base population for our selective breeding experiment and as the source population for our iso-female inbred lines for our genome-wide phenotypic association study. This line was created from wild caught wasps collected from Hoge Veluwe National Park in the Netherlands and is maintained as large outbred population at an effective population size of *N*_*e*_>200.

### *The* Nasonia vitripennis *Genome Reference Panel 1.0*

#### NGRP lines

Iso-female lines were established by randomly collecting 105 virgin females from the HVRx outbred population at HVRx generation 45 [54], which were individually provided with two hosts for two days at 25°C to produce male offspring. Mothers were stored at 4°C and crossed back to one of her sons (mother-son mating) upon emergence of the male offspring. Mated females were provided with two hosts for two days at 25°C to produce male and female offspring. These lines were inbred by 8 generations of full-sib mating to produce 34 stable inbred lines, followed by random mating. This resulted in an inbreeding coefficient of *F*=0.87, equivalent of 10 generations of diploid full-sib matings.

#### Genome resequencing

DNA was isolated from 60 pooled females at generation 29 for each of the 34 stable inbred lines, using a standard high salt–chloroform protocol [102]. Library construction and genome sequencing were performed by BGI Tech Solutions according to standard Illumina protocols. For each inbred line, a 91bp paired-end library was constructed and the libraries were run on an Illumina HiSeq 2000 (Illumina, San Diego, CA, USA). Clean reads were aligned to the *Nasonia vitripennis* genome build Nvit_2.1 (GCF_000002325.3, downloaded from the NCBI FTP site: ftp://ftp.ncbi.nlm.nih.gov/genomes/all/GCF/000/002/325/GCF_000002325.3_Nvit_2.1) using BWA v0.6.2-r126 [103]. Duplicated reads were filtered out using samtools v0.1.18 [104] and alignments were generated as BAM files for further analysis. Assembled data have been submitted to the NCBI Short Read Archive in BioProject no. PRJNA387118 and with accession no. SRP107298. SNPs were called for each line simultaneously using the Joint Genotyper for Inbred Lines (JGIL) v1.6 [55] using the following parameters: read mapping quality threshold: 10, number of generations: 10. SNPs with a quality less than 20 were ignored, and the genotypes of any individual for which the SNP quality score was less than 10 were treated as missing genotypes. SNP data were stored as VCF and have been deposited in the EMBL European Variation Archive in Project no. PRJEB33514 with analysis accession no. ERZ1029220.

#### Population genomics

For the estimation of nucleotide diversity (*π*) and Tajima’s *D*, BAM files resulting from the above described analysis were merged (samtools merge) separately for the HVRx outbred population bam files and the inbred line bam files to produce one outbred BAM file and one inbred bam file. BAM files were combined without a priori coverage correction. From these BAM files, pileup files were produced (samtools mpileup) and subsampled to standardize the coverage at 20X using subsample-pileup.pl of PoPoolation [105] and the following settings target-coverage 20, max-coverage 400, min-qual 20, method withreplace. These subsampled pileup files, were used to estimate the nucleotide diversity (*π*) and Tajima’s *D* in sliding windows using variance-sliding.pl of PoPoolation [105] and the following settings pool-size 60, min-count 5, min-coverage 10, min-covered-fraction 0.5, fastq-type sanger, window-size 400000, step-size 200000.

Linkage disequilibrium (*r*^2^) was estimated between all pairs of SNPs within a distance of 1 MB using the VCF resulting from the JGIL SNP calling pipeline of the inbred lines as input. To estimate LD decay, we fitted the equation *r*^2^ = 1/(1+px) for every focal SNP (using nls method, [106] where x denotes the distance to every other SNP 1MB up- and 1MB down-stream of the focal SNP. After the value for p was retrieved for every SNP, the distance at which LD equal 0.5 was estimated by (1/p). Then these values were plotted against the position of the SNPs to get an indication of LD variation in the genome. To determine the need for correction based on population structure, we determined the significance of the correlation between pairwise *F*_*ST*_ values over all SNPs called in the JGIL pipeline and pairwise absolute Euclidian distance in sex ratio for the 26 lines included in the GWAS (see below) using Spearman’s rank correlation. When correlated with the phenotype, population structure can result in false positives in GWAS analyses, requiring a correction based on the relatedness of the samples [107].

### Wolbachia *detection*

*Nasonia vitripennis* can be infected with the maternally inherited bacterium *Wolbachia pipentis*, which affects reproduction. To be able to account for the effects of *Wolbachia*, the NGRP lines were assessed for *Wolbachia* infection in generation 31 after initial inbreeding using a PCR assay for the *Wolbachia* specific *wsp* gene (81F/691R primers; [108]. PCR conditions were denaturation at 94 °C for 3 min, then 35 cycles of 94 °C for 1 min, 55 °C for 1 min, 72 °C for 1 min and a final extension at 72 °C for 5 min. PCR products were visualized on a 1% agarose gel stained with ethidium bromide.

#### Iso-female line GWAS

The focal females used in this experiment were from 26 *Wolbachia*-positive NVGRP lines. At the point of phenotyping associated with this experiment, these lines had been mass-reared for 36 generations after initial inbreeding.

To control for possible host and other maternal effects, we isolated 40 females (2 day old, mated) from the mass culture of each of the 26 lines into individual glass vials and provided each with three hosts. We then used 40 females from the resulting F1 generation in the experiment, one female per “grandmother”, per line. These experimental females (2 day old, mated) were isolated into individual glass vials and provided with a single host for 24 hours as a pre-treatment to facilitate egg development. Pre-treatment hosts were discarded, and each female given a piece of filter paper soaked in honey solution for a further 24 hours. Subsequently, we gave experimental females access to three hosts for a period of 24 hours. One-way escape tubes were fitted to the glass vials after 1-hour had passed to allow females to disperse, preventing unnatural levels of superparasitism. Hosts were incubated and we counted and sexed the emerging offspring to calculate sex ratio. In total, we phenotyped 57,465 wasps in 958 broods (mean clutch size=60.04).

We analysed the variation in sex ratio in two steps. First, we fitted a Generalised Linear Model with binomial error structure and a logit link function to test for significant differences in sex ratio between the iso-female lines. To correct for over-dispersion, common when analysing binomial data, *F*-tests were used. Second, we determined the between iso-female line variation using linear mixed-effect models on arcsine square root transformed data for sex ratio, and untransformed data for clutch size. Iso-female line was fitted as a random effect and variance components were estimated by REML. This isofemale line analysis estimates the genetic variation as the broad-sense heritability *H*^2^ [51, 109–111]. All statistical analyses were carried out in R [106].

#### Genotype-phenotype associations

To identify SNPs significantly associated with single-foundress sex ratio, we performed single marker regressions for all SNPs on each chromosome segregating between iso-female lines using a general linear mixed effect model. SNPs segregating within lines were treated as missing data. SNPs were excluded if the allele count was less than 3. A Generalized Linear Mixed Model was implemented using the glmer function in the lme4 package [112] in R [106], with sex ratio as the response variable, genotype as the explanatory variable, and line as a random effect. We used a binomial error structure and a logit link function. *P*-values for the significance of the association between single-foundress sex ratio and each SNP were obtained. Significance thresholds were estimated by permutating iso-female line identity for each SNP. The permutated dataset was then used to estimate an empirical threshold using a q-value of 0.1 as significant, which resulted in an empirical threshold of *P*=2.00^*^10^−6^, or – *log*_*10*_*(P)*= 5.70 for sex ratio.

Similar single marker regressions using a linear mixed model were performed for clutch size, using the lmer function in the lme4 package [112] in R [106], with clutch size as the response variable, genotype as the explanatory variable, and line as a random effect. To determine significance thresholds for clutch size, we permutated iso-female line identity for each SNP. Because the minimum q-value was 0.463, no variant was considered significant for clutch size.

To determine the scope for a shared genetic background, we determined the overlap in test statistics between sex ratio and brood size in 400 kb windows per chromosome. To do this, we calculated the overlap in SNPs for an increasing rank in p-values. We then performed he SuperExact test in the SuperExactTest package [113] in R on this overlap for each cutoff obtained observed p-values. Because genome structure can have an effect on the result, mainly through variation in the number of SNPs in each window, we also performed the same analyses on 100 randomized sites where SNP identity was permutated. Windows were considered significant if the observed p-value was lower than any of the 100 permutated ones. We used an increasing windows size of 25, 50, 100, 200 and 400 KB windows (Supplementary Figure 4).

#### Identification of candidate genes

To identify candidate genes, we annotated the VCF file with SnpEff v4.3b [114] based on the NCBI Nasonia vitripennis Annotation Release 102 (https://www.ncbi.nlm.nih.gov/genome/annotation_euk/Nasonia_vitripennis/102/). Candidate SNP positions and their effects were filtered from the annotated VCFs using SnpSift v4.3b [114].

## Declarations

## Acknowledgements

We would like to thank Gabriella Bukovinszkine Kiss, Bertha Koopmanschap and Jade Green for their help in the laboratory, and Eveline Verhulst for her insightful comments on the molecular biology of sex determination.

## Availability of data and material

Raw DNA sequencing reads deposited in NCBI with the project accession of PRJNA387118. SNP data were stored as VCF and have been deposited in the EMBL European Variation Archive (EVA) in Project no. PRJEB33514 with analysis accession no. ERZ1029220. The NVGRP iso-female lines are available upon request from the lab of Bart Pannebakker

## Authors’ contributions

BAP, LvdZ and DMS conceived the study. NC performed the sex ratio experiments, BAP, NC and JvdH perfomed the statistical and sequence analyses. BAP and DMS wrote the manuscript. All authors read and approved the final manuscript.

## Consent for publication

Not applicable.

## Ethics approval and consent to participate

Not applicable.

