## Supplementary Material for "Genomics of sex allocation in the parasitoid wasp *Nasonia vitripennis*"

**Supplementary Tables**

Supplementary Table 1 Sequencing statistics NVGRP. For each NVGRP iso-female line, the total number of clean reads (million reads), the total number of clean bases (Gb), percentage of nucleotides with a read quality higher than 20 (Q20 percentage), GC content (GC %), total number of reads mapped to Nvit 2.1 assembly (million reads), total number of nucleotides covered (Mb) and the percentage of Nvit 2.1 assembly covered (percentage) is given. Data in SupplementaryTable1.xlsx

Supplementary Table 2 NVGRP Sliding windows analysis (400kb windows). Data in SupplementaryTable2_PopoolationResults.xlsx

Supplementary Table 3 Results for sex ratio and clutch size GWAS in NVGRP. Data in SupplementaryTable3_GWAS_Candidate_and_All_SNPs.xlsx

Supplementary Table 4 Linkage disequilibrium ($r^2$) between 18 SNPs significantly associated with sex ratio in the 25 NVGRP lines

| | NC_015867.2_30565627 | NC_015867.2_30570243 | NC_015867.2_30593991 | NC_015867.2_30596018 | NC_015867.2_30598225 | NC_015867.2_30602783 | NC_015867.2_30602789 | NC_015867.2_30604770 | NC_015867.2_30606952 | NC_015867.2_30607468 | NC_015867.2_30608905 | NC_015867.2_30608907 | NC_015867.2_30610368 | NC_015867.2_30612505 | NC_015867.2_30643961 | NC_015867.2_30670045 | NC_015867.2_30682477 | NC_015867.2_31141049 |
|---|---|---|---|---|---|---|---|---|---|---|---|---|---|---|---|---|---|---|
| **NC_015867.2_30565627** | 1.000 | | | | | | | | | | | | | | | | | |
| **NC_015867.2_30570243** | 1.000 | 1.000 | | | | | | | | | | | | | | | | |
| **NC_015867.2_30593991** | 0.909 | 0.909 | 1.000 | | | | | | | | | | | | | | | |
| **NC_015867.2_30596018** | 0.909 | 0.909 | 1.000 | 1.000 | | | | | | | | | | | | | | |
| **NC_015867.2_30598225** | 0.909 | 0.909 | 1.000 | 1.000 | 1.000 | | | | | | | | | | | | | |
| **NC_015867.2_30602783** | 0.909 | 0.909 | 1.000 | 1.000 | 1.000 | 1.000 | | | | | | | | | | | | |
| **NC_015867.2_30602789** | 0.909 | 0.909 | 1.000 | 1.000 | 1.000 | 1.000 | 1.000 | | | | | | | | | | | |
| **NC_015867.2_30604770** | 0.909 | 0.909 | 1.000 | 1.000 | 1.000 | 1.000 | 1.000 | 1.000 | | | | | | | | | | |
| **NC_015867.2_30606952** | 0.909 | 0.909 | 1.000 | 1.000 | 1.000 | 1.000 | 1.000 | 1.000 | 1.000 | | | | | | | | | |
| **NC_015867.2_30607468** | 0.909 | 0.909 | 1.000 | 1.000 | 1.000 | 1.000 | 1.000 | 1.000 | 1.000 | 1.000 | | | | | | | | |
| **NC_015867.2_30608905** | 0.909 | 0.909 | 1.000 | 1.000 | 1.000 | 1.000 | 1.000 | 1.000 | 1.000 | 1.000 | 1.000 | | | | | | | |
| **NC_015867.2_30608907** | 0.909 | 0.909 | 1.000 | 1.000 | 1.000 | 1.000 | 1.000 | 1.000 | 1.000 | 1.000 | 1.000 | 1.000 | | | | | | |
| **NC_015867.2_30610368** | 0.909 | 0.909 | 1.000 | 1.000 | 1.000 | 1.000 | 1.000 | 1.000 | 1.000 | 1.000 | 1.000 | 1.000 | 1.000 | | | | | |
| **NC_015867.2_30612505** | 0.909 | 0.909 | 1.000 | 1.000 | 1.000 | 1.000 | 1.000 | 1.000 | 1.000 | 1.000 | 1.000 | 1.000 | 1.000 | 1.000 | | | | |
| **NC_015867.2_30643961** | 0.909 | 0.909 | 1.000 | 1.000 | 1.000 | 1.000 | 1.000 | 1.000 | 1.000 | 1.000 | 1.000 | 1.000 | 1.000 | 1.000 | 1.000 | | | |
| **NC_015867.2_30670045** | 0.909 | 0.909 | 1.000 | 1.000 | 1.000 | 1.000 | 1.000 | 1.000 | 1.000 | 1.000 | 1.000 | 1.000 | 1.000 | 1.000 | 1.000 | 1.000 | | |
| **NC_015867.2_30682477** | 0.909 | 0.909 | 1.000 | 1.000 | 1.000 | 1.000 | 1.000 | 1.000 | 1.000 | 1.000 | 1.000 | 1.000 | 1.000 | 1.000 | 1.000 | 1.000 | 1.000 | |
| **NC_015867.2_31141049** | 0.822 | 0.822 | 0.909 | 0.910 | 0.910 | 0.904 | 0.904 | 0.904 | 0.904 | 0.904 | 0.909 | 0.909 | 0.904 | 0.904 | 0.904 | 0.904 | 0.904 | 1.000 |

### Supplementary Figures

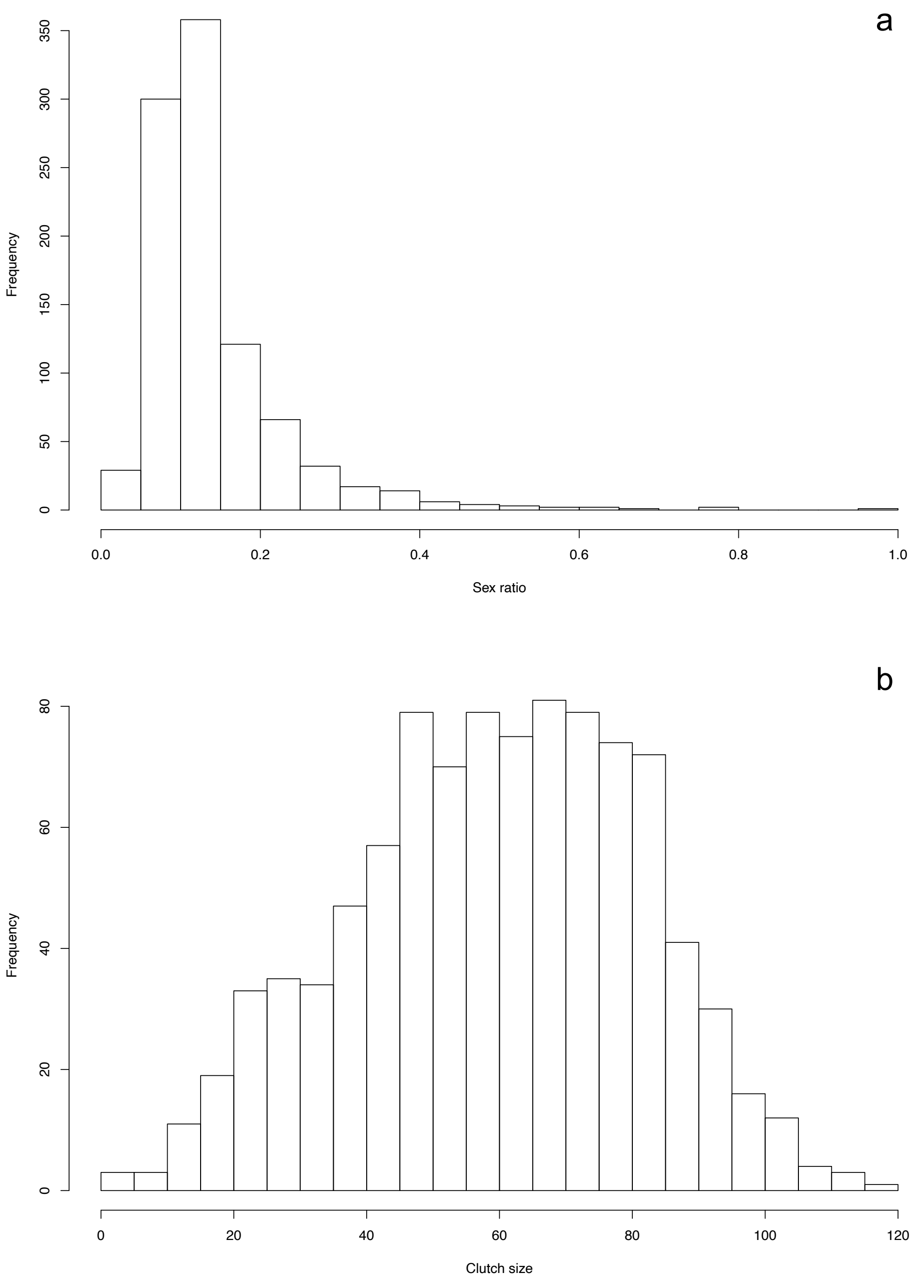


*Supplementary Figure 1.* Distribution of phenotypic values over all 26 tested NVGRP lines, for (a) sex ratio expressed as proportion males, and (b) clutch size.

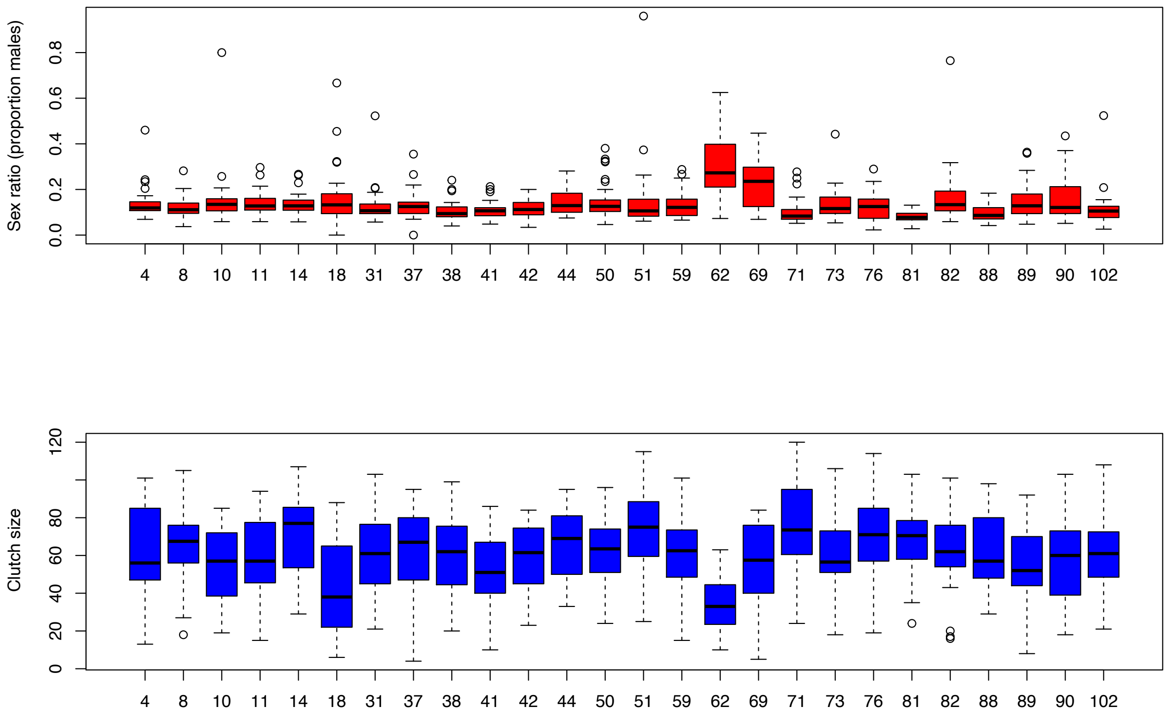


*Supplementary Figure 2.* Boxplot of sex ratio (proportion males) and clutch size among all 26 tested NVGRP lines. Plot shows the median, the box indicates the first and the third quartile, the whiskers indicate 1.5 times the interquartile range and values beyond that are labelled outliers (open circles).

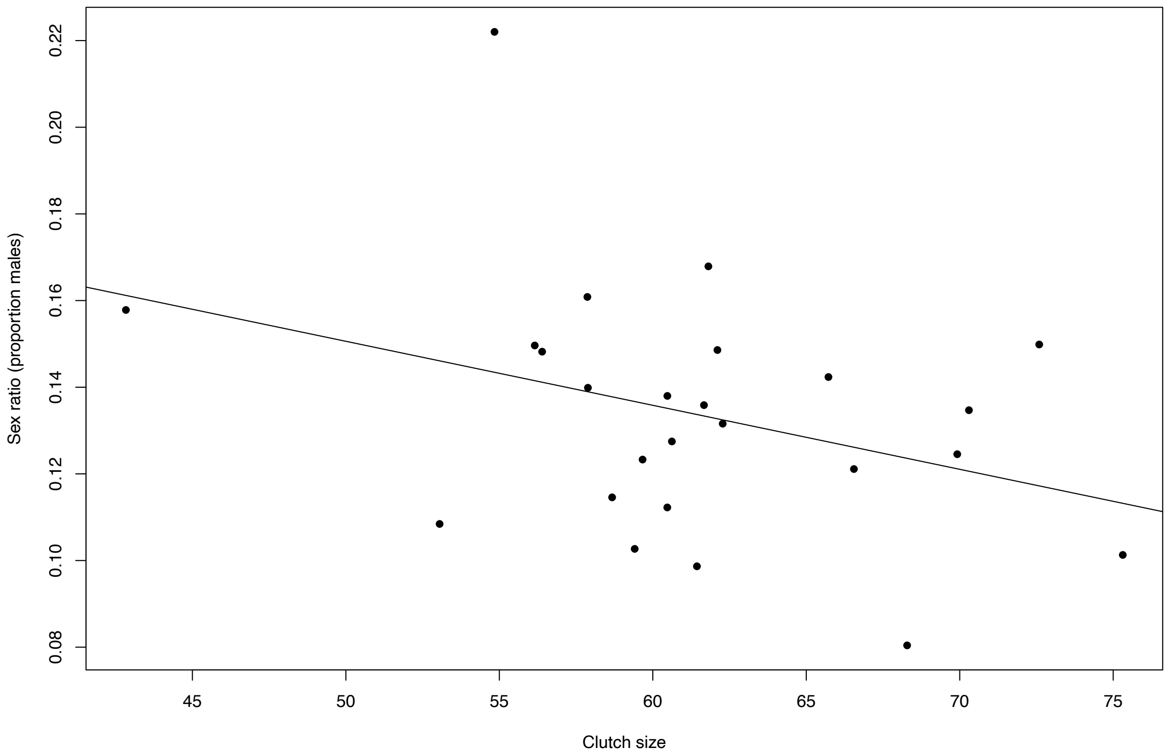


*Supplementary Figure 3.* Scatterplot of the mean clutch size and the mean sex ratio for the 25 tested NVGRP lines (excluding outlier line 62). Solid line shows the fit of the linear regression y= 0.224467 -0.001477x.

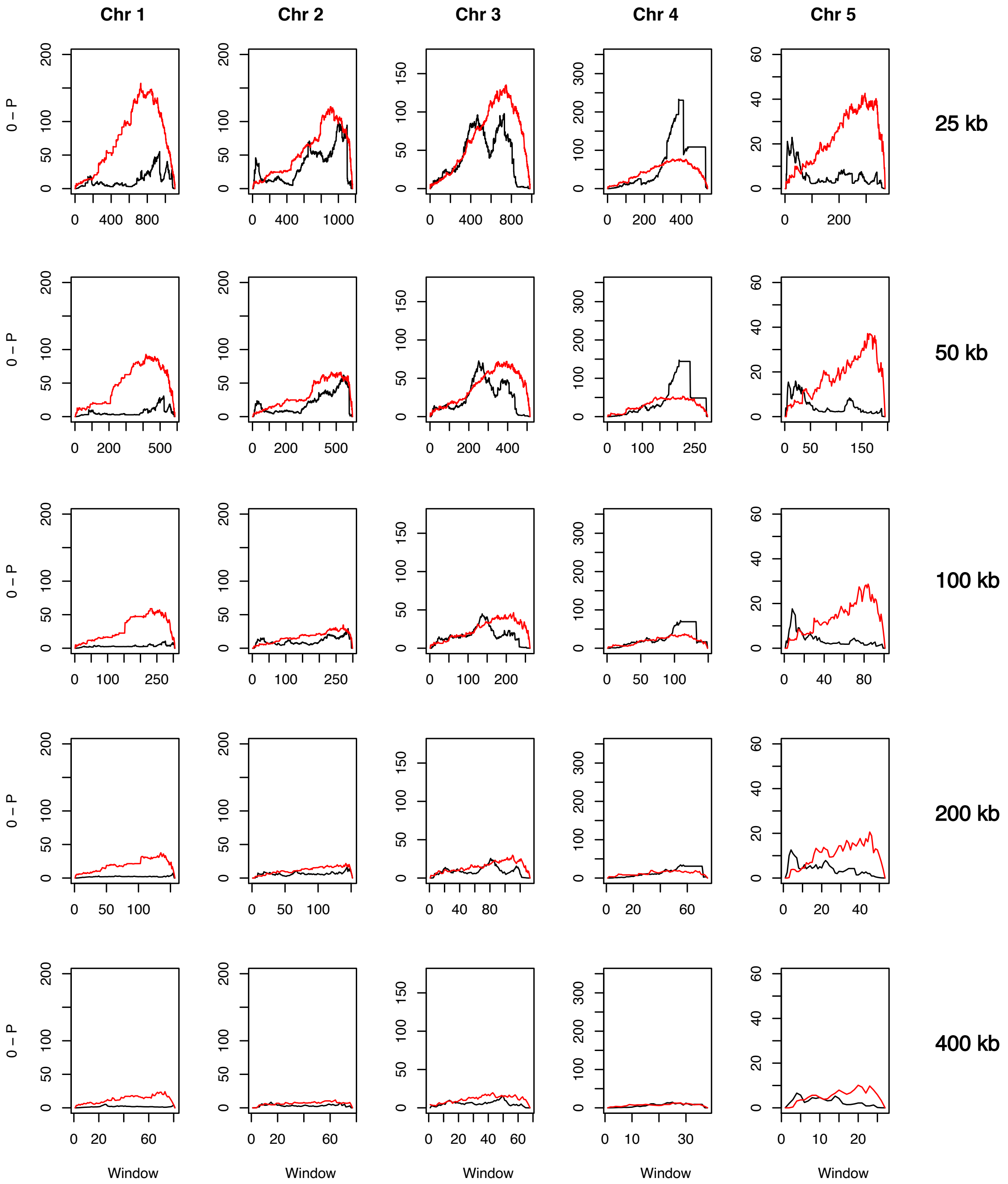


*Supplementary Figure 4.* Multipaneled plot showing the cumulative results of the SuperExactTest for association of the p-values of the sex ratio and clutch size GWAS per chromosome for windows of 25kb, 50kb, 100kb, 200kb and 400kb. Black solid line shows observed overlap using GWAS p values for each SNPs, red solid line shows the most significant p-value for each window given from a distribution of 100 permutations. Significant overlap is observed for all window sizes, when the p-values of the association between the sex ratio and clutch size GWAS exceeds the p-values of the permutation.
